# Monitoring Neural Activity During Exposure to Drugs of Abuse With In Vivo Fiber Photometry

**DOI:** 10.1101/487546

**Authors:** Jennifer A. Rinker, Dominic Gioia, Kevin M. Braunscheidel, Wesley N. Wayman, Michaela Hoffman, Linsey Passarella, Erin S. Calipari, Patrick J. Mulholland, John J. Woodward

**Author notes:** To Whom Correspondence Should Be Addressed, John J. Woodward, PhD, Department of Neuroscience, Medical University of South Carolina, Charleston, SC 29425.

## Abstract

Drugs of abuse are known to alter activity in areas of brain associated with reward, cognition and decision making. Changes in neural activity in these regions that follow repeated exposures to abused substances may underlie the development of addictive behaviors and contribute to the high rates of relapse associated with drug use. Measuring real-time changes in neural activity during drug seeking and taking is important for correlating changes in behavior with alterations in neuronal signaling typically measured using ex vivo electrophysiological recordings. In this study, C57BL/6J mice or Sprague-Dawley rats were injected in different brain areas with adeno-associated viruses (AAV) encoding the calcium sensor GCaMP6f along with an optical fiber. Calcium-dependent fluorescence was monitored in the nucleus accumbens core or mPFC during and following exposure to toluene vapor and in the medial prefrontal cortex (mPFC) and orbitofrontal cortex (OFC) during ethanol drinking. Toluene vapor, at concentrations previously shown to induce conditioned place preference, produced a rapid decrease in the frequency of calcium transients in the NAc core of rats that recovered following washout of the toluene vapor. In a probabilistic risk task, GCaMP6 signals in rat mPFC increased just prior to lever pressing and showed decreases during the reward phase that were proportional to reward size. Toluene pretreatment elevated the signal during the decision-making period while post-lever responses were independent of reward size. Using the drinking in the dark (DID) protocol in mice, we observed a consistent increase in GCaMP6 fluorescence during the period leading up to an ethanol drinking bout, a decrease during consumption and a rebound increase following the bout. The initial increase in signal prior to consumption was greater for ethanol and sucrose than water. GCaMP6 signals in the lateral OFC also decreased during ethanol consumption and increased following bout completion while no increase in activity was noted prior to bout initiation. Following repeated cycles of chronic intermittent ethanol (CIE) exposure that enhanced ethanol consumption, OFC calcium signals during and after ethanol drinking were similar to those in air-treated animals. Addition of quinine to the ethanol solution augmented the decrease in signal during consumption in both air and CIE mice while having no effect on the magnitude of the rebound in activity. Conversely, when sucrose was added to the ethanol solution, air exposed mice showed blunted changes in GCaMP6 signals while those in CIE mice were enhanced. Overall, the results from these experiments complement and extend data from prior behavioral and electrophysiological studies and support the use of in vivo fiber photometry in the study of effects of abused substances on brain function.

## Introduction

Understanding how drugs of abuse affect neural circuits involved in initiating and controlling motivated behaviors is an important goal of addiction neuroscience research. Historically, studies investigating these effects have largely relied upon rodent models of drug seeking and taking followed by ex vivo slice electrophysiology recordings that provides a snapshot of drug-induced changes in neuronal excitability and synaptic signaling. While informative, these approaches suffer from the lack of measurements of neural activity that is time-locked to behaviors associated with drug seeking and taking. In vivo electrophysiological recordings circumvent some of these limitations but lack the ability to easily and reliably identify sub-populations of neurons, presynaptic activity or signaling of specific drug-related circuits. In addition, although in vivo 2-photon microscopy and mini-endoscopic imaging approaches offer online, high-resolution imaging, their use may be limited by imaging depth, use of head-fixed animals, damage due to implantation of large diameter lenses and often significant up-front equipment and maintenance costs.

An alternative to these approaches is the use of in vivo fiber photometry that uses off the shelf optical components and genetically encoded calcium sensors such as GCaMP to track activity in defined neural circuits in freely behaving animals (Siciliano and Tye, 2018). These approaches were initially developed to study neural mechanisms and cell types implicated in mediating various aspects of neuropsychiatric and neurodegenerative disorders (Gunaydin *et al*, 2014; Lerner *et al*, 2015) and are now increasingly being applied to study of various drugs of abuse. For example, Calipari and colleagues used GCaMP6f fiber photometry to show that D1 medium spiny neurons in the nucleus accumbens (NAc) encode information required to induce a conditioned place preference (CPP) to cocaine (Calipari *et al*, 2016). A subsequent study by this group also using fiber photometry showed that cocaine’s effect on calcium transients in the ventral tegmental area was greater in estrus-stage female mice as compared to males or di-estrus females (Calipari *et al*, 2017).

These findings and those from studies discussed below have established in vivo fiber photometry as a viable approach for studying how a wide range of drugs of abuse alter the function of neurons in key nodes of the addiction neurocircuitry of the brain.

## Methods

### Animals

Male C57BL/6J mice were obtained from Jackson Laboratories (Bar Harbor, ME) (https://www.jax.org/strain/013636) and were at least 7 weeks of age upon arrival. They were group-housed (4/cage) and allowed to acclimatize to the colony room for at least one week in a temperature and humidity controlled AAALAC-approved facility. Male Sprague-Dawley rats were purchased at 21 days of age from Harlan Laboratories and pair-housed. Mice and rats were maintained on a 12-hour light/dark cycle with lights off at 09:00 am and had ad libitum access to food and water except where noted. All animals were treated in strict accordance with the NIH Guide for the Care and Use of Laboratory Animals and all experimental methods were approved by the Medical University of South Carolina’s Institutional Animal Care and Use committee.

### Viral Injection and Fiber Surgeries

Adult male C57BL/6J mice and Sprague-Dawley rats underwent stereotaxic surgery for viral injection and ferrule implantation 4-6 weeks prior to the start of photometry recordings. Animals were anesthetized using isoflurane mixed with medical-grade air (5% induction, 1-2% maintenance), given a subcutaneous injection of 5mg/kg carprofen (Pfizer), and placed in a Kopf stereotaxic instrument with digital display. Adeno-associated virus (AAV) encoding GCaMP6f under the control of a synapsin or CaMKII promoter (Addgene) was injected into target areas in isoflurane-anesthetized animals. Coordinates for viral injection (in mm; relative to bregma) were rat mPFC (AP +2.95, lateral ± -0.60, ventral -2.85); rat NAc core (AP +2.0, lateral ± 1.4, ventral -6.0); mouse lateral OFC (AP +2.46, lateral ± 1.3, ventral -2.4); medial PFC (AP +1.70, lateral ± 0.40, ventral -2.35). For each region, virus was unilaterally infused at a rate of 100 nL/min for (200 nL for mouse mPFC, OFC; 300 nL for rat mPFC; 150 nL for rat NAc) and microinjection needles were left in place for 5 min post-infusion. Following viral infusion, custom-made optical fibers (Thorlabs, 400 µm, 0.48 NA) connected to stainless steel or ceramic ferrules (Thorlabs, outer diameter 1.25 mm mouse, 2.5 mm rat) were implanted. Following the conclusion of behavioral experiments, rodents were transcardially perfused with Dulbecco’s PBS and 4% paraformaldehyde and ferrule placements were histologically verified via the visualization of fiber tracks and viral vector transduction efficiency via GCaMP6f expression.

### Toluene Vapor Exposure

Rats were exposed to toluene vapor as described previously (Beckley *et al*, 2013). Briefly, animals were placed in a custom-made, flow-through chamber and connected to an optical fiber attached to optical swivel (FRJ_1x1; Doric Lenses) mounted on the top of the chamber. The chamber was continuously perfused with air (5 liters/min) and at discrete times, the airflow was diverted through dual sevoflurane vaporizers filled with toluene that provided toluene vapor concentrations up to 10,500 ppm followed by washout of the toluene vapor. Vapor concentrations were measured in real-time using a toluene sensor Fiber photometry data were collected continuously throughout the baseline, toluene and washout periods.

### Probabilistic Discounting

An automated probabilistic discounting setup was used to assess risky decision making in Sprague Dawley Rats (St Onge and Floresco, 2010). Briefly, rats aged P60 were food restricted to 85-90% of their free feeding weight and trained to lever press to receive a palatable food reward (20% sweetened condensed milk) on a FR1 schedule. During each 1 h session, one lever delivered a small, certain reward (30 µl, 100% of the time), while a second lever delivered a large, uncertain reward (90 µl, reinforcement probability varied between 5 successive blocks of 18 trials: 8 forced choice, 10 free choice). Between each block, the odds of risky reinforcement descended from 100% to 6.25%. The primary dependent measure for this task is the proportion of risky lever choices made during free choice trials in each of these five risk blocks. Testing began after choice preference in the first block was > 80% and stable responding was achieved: two-way ANOVA on three consecutive testing days yields no block x day interaction or main effect of day (p >0.1). One day prior to photometry recordings, rats were habituated in an identical operant conditioning chamber outfitted for fiber photometry recordings. For the next two days, rats were exposed to either 10,5000 ppm toluene or air for 15 minutes, returned to their homecage for 30 minutes, and then recorded during probabilistic discounting.

### Drinking in the Dark

Water, ethanol, and sucrose consumption was induced using a standard 4-day drinking in the dark (DID) protocol (Rhodes *et al*, 2005; Rinker *et al*, 2017). Approximately 3 hours into the dark cycle, home cage water bottles were removed, and animals were given access to a bottle containing water, 20% v/v ethanol, or 10% w/v sucrose for 2 hours on days 1–4. Fiber photometry recordings were performed on day 4 of drinking. Mice had access to water during test week 1 and ethanol during test weeks 2-4. Following a two- to four-week abstinence period, mice were then given access to sucrose in their home cages.

### Operant Ethanol Self-Administration

Mice were trained to self-administer 10% ethanol under an FR1 schedule of reinforcement using a post-prandial drinking schedule. During initial magazine and lever training, mice were food deprived for 23 hours a day and given access to food during the 1 hour before each self-administration session. Mice were given ad libitum access to water except for the 1 hour feeding period prior to self-administration. Mice maintained normal bodyweights during this procedure. During the first two weeks, mice underwent magazine training and were given non-contingent access to 10% ethanol in Med Associates boxes during 1 hr sessions. After magazine training, mice went through 8 weeks of ethanol self-administration under an FR1 schedule. When mice pressed the active lever, house lights turned off for 1 second, a tone was presented and ethanol was delivered to a drinking well in the operant box; while, nothing happened when the inactive lever was pressed.

### Chronic Intermittent Ethanol (CIE) Exposure

Mice were treated with repeated cycles of ethanol vapor exposure as previously described (Nimitvilai *et al*, 2016). Briefly, each cycle consisted of daily exposure to ethanol vapor for 16 hr followed by 8 hr of abstinence in the home cage. This was repeated each day for 4 consecutive days followed by three days of abstinence before beginning the next cycle of ethanol exposure. Ethanol (95%) was volatized by passing air through a submerged air stone and the resulting vapor was mixed with fresh air and delivered to Plexiglas inhalation chambers (5 L/min) to maintain consistent ethanol concentrations between 17-21 mg/L air in the chamber. This yielded blood ethanol concentrations (BEC) in the range of 150-250 mg/dl. Prior to entry into the ethanol chambers, CIE mice were injected intraperitoneally (20 ml/kg body weight) with ethanol (1.6 g/kg; 8% w/v) and the alcohol dehydrogenase inhibitor pyrazole (1mmol/kg) to maintain stable BECs. Control mice were injected with saline and pyrazole before being placed in air chambers. Chamber ethanol concentrations were monitored daily and air flow was adjusted to maintain concentrations within the specified range. In addition, blood samples were collected and determined from all animals to monitor BECs during the course of inhalation exposure.

### In Vivo Fiber Photometry Imaging

Data were acquired using custom-built imaging equipment based on that described by the Deisseroth lab (Lerner *et al*, 2015) with modifications. Illumination was provided by 405 nm and 490 nm fiber collimated LEDs (Thorlabs; 30 µW per channel for OFC/NAc; 15 µW for mPFC) connected to a four-port fluorescence mini-cube (Doric Lenses). The combined LED output passed through a 400 µm optical fiber (0.48 NA) pigtailed to a rotary joint (FRJ_1x1_PT, Doric Lenses) and connected to the implanted fiber using a ceramic sleeve or pinch connector (Thorlabs). Emission light was focused onto a photodetector (Newport model 2151; DC low setting) low-passed filtered at 3 Hz and sampled at 6.1 kHz by a RZ5P lock-in digital processor (TDT) controlled by Synapse software (TDT). Excitation light was sinusoidally modulated at 531 Hz (405 nm) and 211 Hz (490 nm) via software control of an LED light driver (Thorlabs). Real-time demodulated emission signals from the two channels were acquired at a frequency of 0.93084 kHz and stored offline for analysis. In the drinking studies, data from lickometers, head entry sensors and lever presses (MedAssociates) were also collected via TTL inputs to the digital processor.

### Data Analysis

Data were processed using custom written functions in Matlab (Mathworks) software and analyzed for statistical significance using Prism (Graphpad). Because the 405 nm and 490 nm signals showed differences in photobleaching over time (slope measured over single recordings sessions, mean ± sem; 405 nm -0.056 ± 0.001, 490 nm -0.0042 ± 0.0004, N=45), the signals for each channel were first fitted to a polynomial versus time and then subtracted from one another to calculate the ΔF/F time series. In mPFC recordings, the mean of the negative values was first subtracted from the normalized Δ*F*/*F* to establish a new baseline of the recordings for comparing changes during ethanol consumption.

For detection and quantitation of transients, the ‘findpeaks’ function in Matlab was used with values for minimum peak prominence, minimum and maximum peak widths and minimum peak distance selected by iterative analysis of the data. Transients with a peak amplitude below a threshold of 2X the median absolute deviation of the normalized signal were excluded from the analysis (Calipari et al., 2016, PNAS). To analyze calcium signals during drinking sessions, lickometer TTL pulses generated by MedAssociates software were used to define bout start times and duration and ΔF/F values were time-locked to episodes (before, during and after) surrounding each bout.

## Results and Discussion

### Toluene and NAc Calcium Transients

Toluene is the prototypical abused inhalant and is a major ingredient of a wide variety of industrial and household solvents. Inhalants such as toluene are often the first drug of abuse tried by children and adolescents and as compared to use of illicit drugs by these individuals it is second only to THC (Johnston *et al*, 2013). Work from the Woodward laboratory has shown that toluene has actions on a wide variety of ion channels that regulate neuronal activity and can induce selective changes in signaling properties of mPFC and VTA DA neurons that project to the NAc (Beckley *et al*, 2013; Wayman and Woodward, 2018b). To begin to understand how toluene vapor affects NAc neuron activity in vivo, we expressed GCaMP6f in the NAc core of adolescent Sprague-Dawley rats and monitored activity using in vivo fiber photometry during toluene vapor exposure. As shown in Figure 1, NAc neurons show robust and frequent calcium transients under baseline conditions. These transients subside when the concentration of toluene vapor was ramped from 0-3,000 ppm and then quickly recover following the washout period. Analysis of the ΔF/F signal shows that toluene vapor significantly reduced the frequency of NAc calcium transients while there was no statistically significant effect on amplitude or width. This may reflect the smaller number of episodes detected during the exposure period. We also noted that the intensity of the overall GCaMP6f signal during toluene exposure decreased as compared to pre- and post-exposure consistent with reduced activity of NAc neurons.

**Figure 1.**
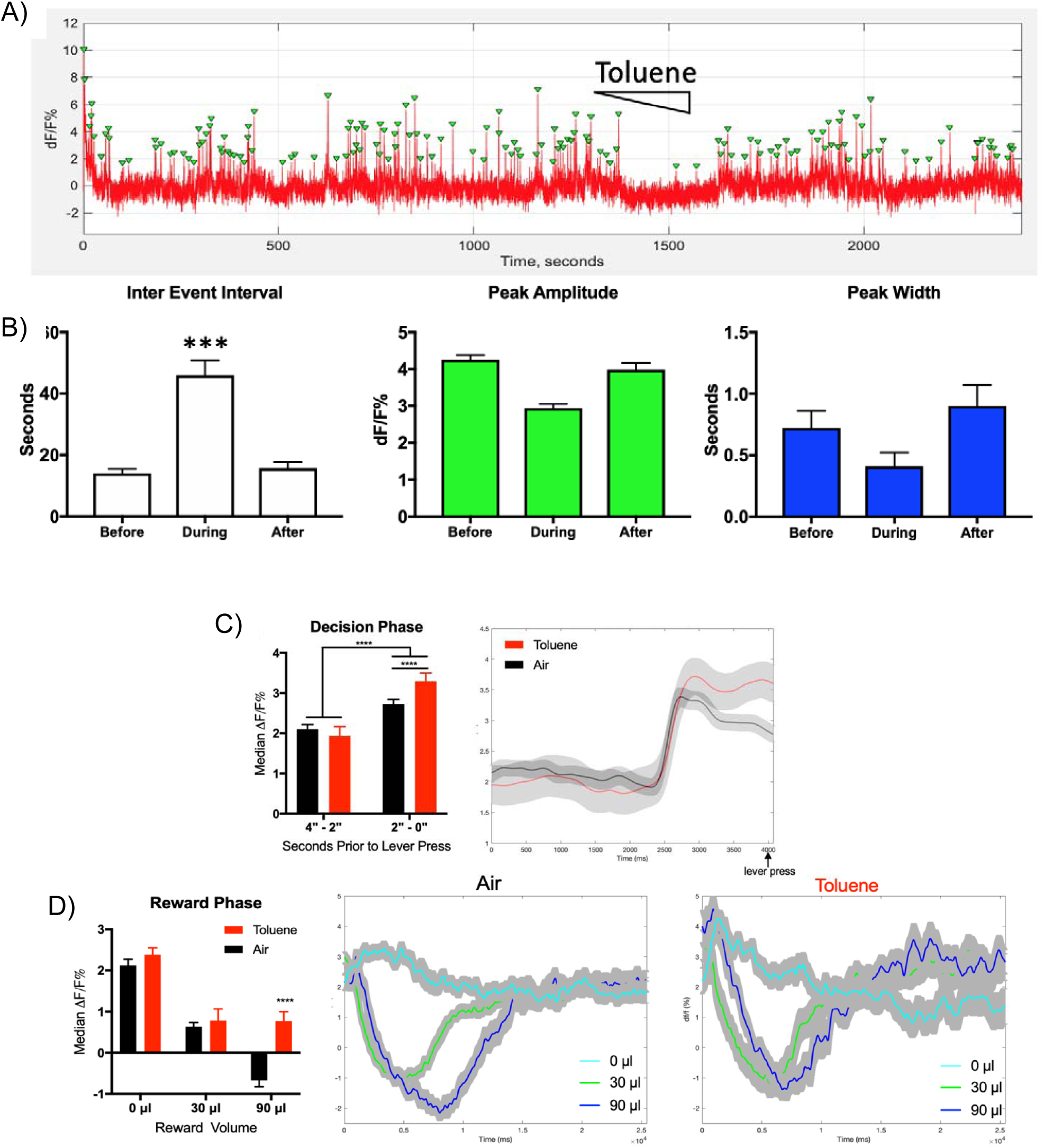
Effect of toluene vapor exposure on NAc core and mPFC neuronal activity as measured by in vivo fiber photometry. (A) Trace shows the motion-corrected NAc core GCaMP6f signal expressed as ΔF/F% before, during and after exposure to increasing concentrations (0-3,000 ppm) of toluene vapor. Green triangles indicate peaks detected by the findpeaks function of Matlab software. (B) Summary graphs show mean (± SEM) peak frequency (expressed as inter-event interval), amplitude and width during the three epochs. Symbol (***), p < 0.001, 1-way ANOVA. (C) Median ΔF/F% ± SEM values (left panel) beginning 4 sec prior to decision-making period in probabilistic risk task. Representative median ΔF/F% ± SEM trace for one rat across 90 trials (right panel). (D) Median ΔF/F ± SEM during the reward phase (left panel), 15 seconds following lever press with representative traces (right panels). (n=2rats;180 trials; 1- or 2-way Anova main effect and Sidak’s post hoc, ****p<0.0001).

In our previous study (Wayman and Woodward, 2018a), repeated pairings with 3,000 ppm of toluene vapor induced a robust CPP that was accompanied by circuit-selective alterations in the excitability of deep-layer mPFC neurons that project to the nucleus accumbens. In particular, whole-cell slice recordings from CPP expressing animals showed that NAc core projecting neurons from layer 5/6 infralimbic (IL) mPFC neurons displayed enhanced current-evoked spiking while layer 5/6 IL neurons projecting to the NAc shell had reduced excitability. No changes in firing were observed in prelimbic NAc projecting neurons following toluene place conditioning. Data from the fiber photometry studies demonstrating reduced activity in NAc neuron excitability during acute toluene exposure suggests that the combined effects of toluene on mPFC and NAc neurons may underlie the development of toluene CPP. Notably, our previous study also showed that chemogenetic excitation of IL-NAc shell projecting neurons blocked the expression of toluene CPP while inhibition of IL-NAc core projecting neurons had no effect (Wayman *et al*, 2018a). Studies with Cre-dependent rat lines are currently underway to selectively interrogate how toluene vapor affects D1 and D2 MSN activity.

We also used GCaMP6 fiber photometry to measure activity of medial prefrontal cortex (mPFC) neurons in Sprague-Dawley rats trained to lever press for reward (20% sweetened condensed milk) under probabilistic conditions. The GCaMP6f signal was analyzed during two phases: the decision phase, four seconds prior to a lever press; and the reward phase, fifteen seconds following a lever press (Fig 1C) GCaMP6f signaling increased in the seconds leading up to a reinforced lever press similar to that observed in mouse mPFC prior to sucrose or ethanol consumption (Fig 2B). Toluene exposure before task performance caused a significant increase in GCaMP6f signal prior to lever selection (Fig 1C; representative trace, right). During the reward phase, the calcium signal under control conditions was inversely related to the volume of reward delivered. Change in mPFC calcium signal to both reward volumes were temporally similar for the descending portion of the transient. However, the response to a large reward took longer to return to baseline compared to the response to a small reward, suggesting this signal tracks reward consumption (Fig 1D, middle panel). There was also a slight increase in calcium signal observed following an unrewarded trial (i.e. 0 ul delivered) that could reflect loss encoding by the mPFC. Toluene vapor exposure prior to behavioral testing increased the calcium signal specifically during delivery of a large reward (Fig. 1D). The similarity in magnitude and temporal dynamics of GCamp6f transients in response to small versus high reward volumes following pretreatment with toluene vapor could reflect an inability to differentiate between reward magnitude and thus may underlie their poor probabilistic discounting behavior (data not shown).

**Figure 2.**
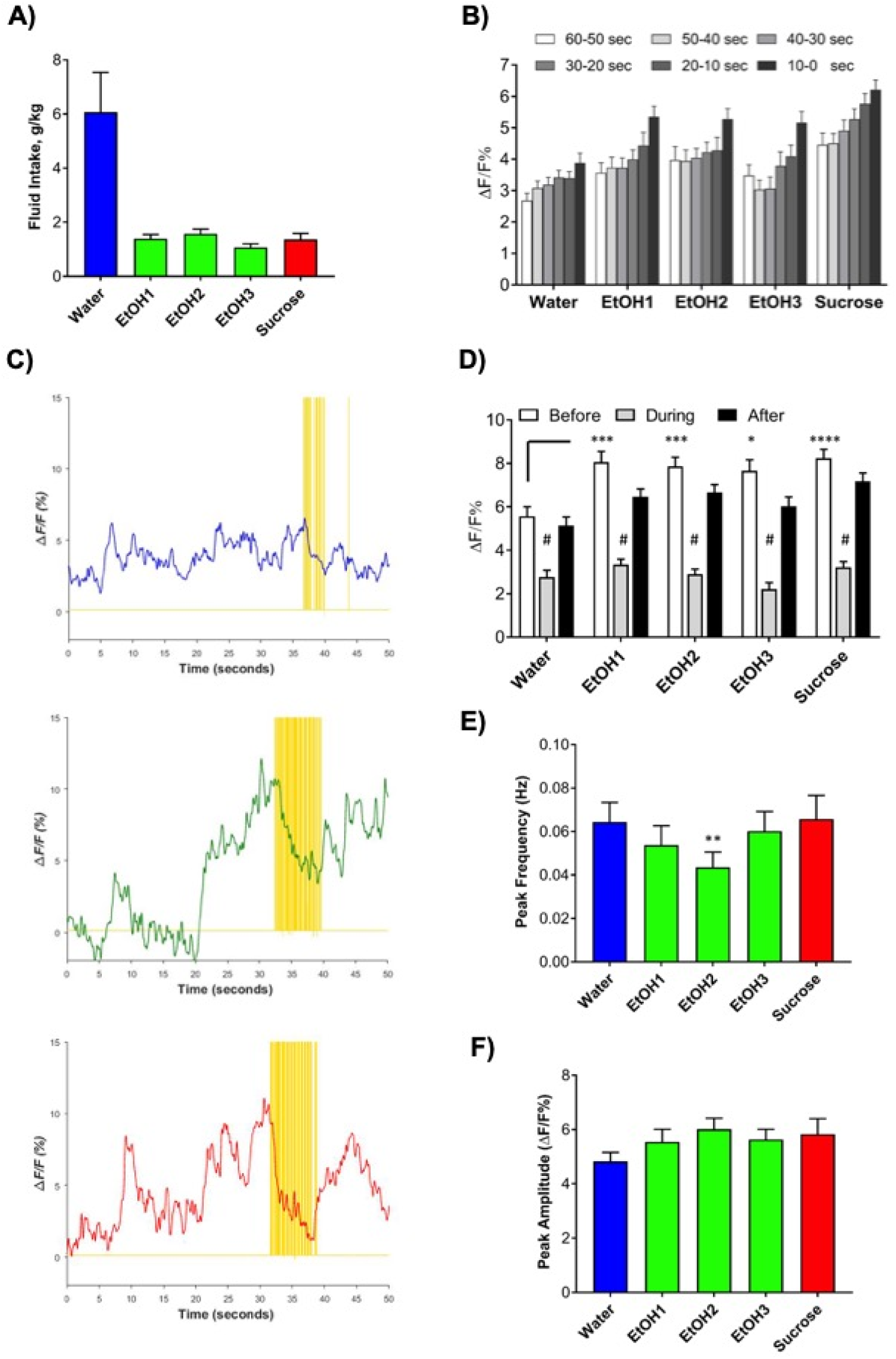
Activity of medial PFC neurons in C57BL/6J mice during water, ethanol and sucrose drinking. (A) Intake of water, ethanol (20%) or sucrose (10%) during drinking in the dark 2-hr sessions (N=9 mice). (B) GCaMP6f signals during sequential 10 sec epochs prior to initiating a bout of drinking. (C) Representative traces of GCaMP6f signaling across 50 seconds surrounding bouts of licking (yellow ticks) for water (top, blue trace), ethanol (middle, green trace) and sucrose (bottom, red trace). D) Averaged changes in GCaMP6f before, during and after initiating drinking. Frequency (E) and amplitude (F) of GCaMP6f transients over the weekly drinking periods. Following the last ethanol drinking session, mice had a 2-4 week withdrawal period prior to sucrose drinking. Data are mean ± sem. Symbols: (*,**,***), p < 0.05, 0.01 0.001; (****, #) p < 0.0001; 1 and 2-way ANOVA.

### Activity of mPFC and OFC neurons during ethanol consumption

In these studies, in vivo fiber photometry was used to monitor activity of neurons in the prelimbic mPFC and lateral OFC of mice during voluntary ethanol consumption. These regions have been implicated in various aspects of alcohol action including reduced control over behavior during acute intoxication and deficits in cognitive function following prolonged use of alcohol (Abernathy *et al*, 2010; Badanich *et al*, 2011). Figure 2 shows results from mPFC studies using the drinking in the dark (DID) model developed to mimic binge-like ethanol consumption in mice. During limited access (2 hr), mice consumed ∼6 g/kg water and ∼1.5 g/kg ethanol and sucrose when tethered to the optical fiber (Fig. 2A). Consistent with previous reports (Horst and Laubach, 2013), the GCaMP6 signal in the prelimbic cortex increased (or ‘ramped’) prior to the initiation of drinking (Fig. 2B). In the vast majority of recorded licking bouts, this signal was well above baseline (>2% Δ*F*/*F*) within 10 seconds of drinking and less than 5% of the drinking bouts had values below baseline within 60 seconds of initiation of licking behavior. Because the peak of the ramped signal occurred immediately before drinking, we quantified the change in GCaMP6 signaling three seconds prior to, during, and three seconds after fluid intake. Prior to the start of intake, there was a peak in the signal that declined during active licking (Fig. 2B, C). A large peak then followed each drinking bout, regardless of the drinking solution. Comparison of the different drinking solutions revealed that the GCaMP6 signal was significantly higher during sessions of ethanol or sucrose drinking as compared to those with water (Fig. 2D). This was most notable for the peak of the ramped signal that preceded ethanol and sucrose drinking bouts versus those for water.

To assess possible change in signal due to toxicity or changes in expression of GCaMP6 over time, we compared the amplitude and frequency of GCaMP6 transients during each drinking session across 7-9 weeks of recordings. The frequency of GCaMP6 peaks was significantly lower in the second week of ethanol drinking compared to peaks during the water drinking session (Fig. 2E). However, the frequency of peaks during access to sucrose return to baseline values. Importantly, the amplitude of GCaMP6 peaks did not differ across the 7-9 weeks of recordings indicating stable expression and recording conditions (Fig. 2F).

Figure 3 shows results from in vivo fiber photometry studies monitoring activity of lateral OFC neurons during operant self-administration of sucrose. A representative example of average and Z-scored ΔF/F values before and after drinking bouts during a 1 hr session of sucrose drinking is shown in Figure 3A (top panels) along with analyses of bout duration, inter-bout interval and licks per bout (Fig. 3A, bottom panels). OFC neurons showed no clear evidence of ramping activity prior to bout initiation but like mPFC neurons, displayed reduced signal during consumption and a rebound increase in activity following bout completion. This is more clearly shown in the summary plot (Fig. 3B) that compares mean ΔF/F values for the 3 seconds prior to bout initiation, the minimum signal obtained during the bout and the maximum signal during the three seconds following bout completion.

**Figure 3.**
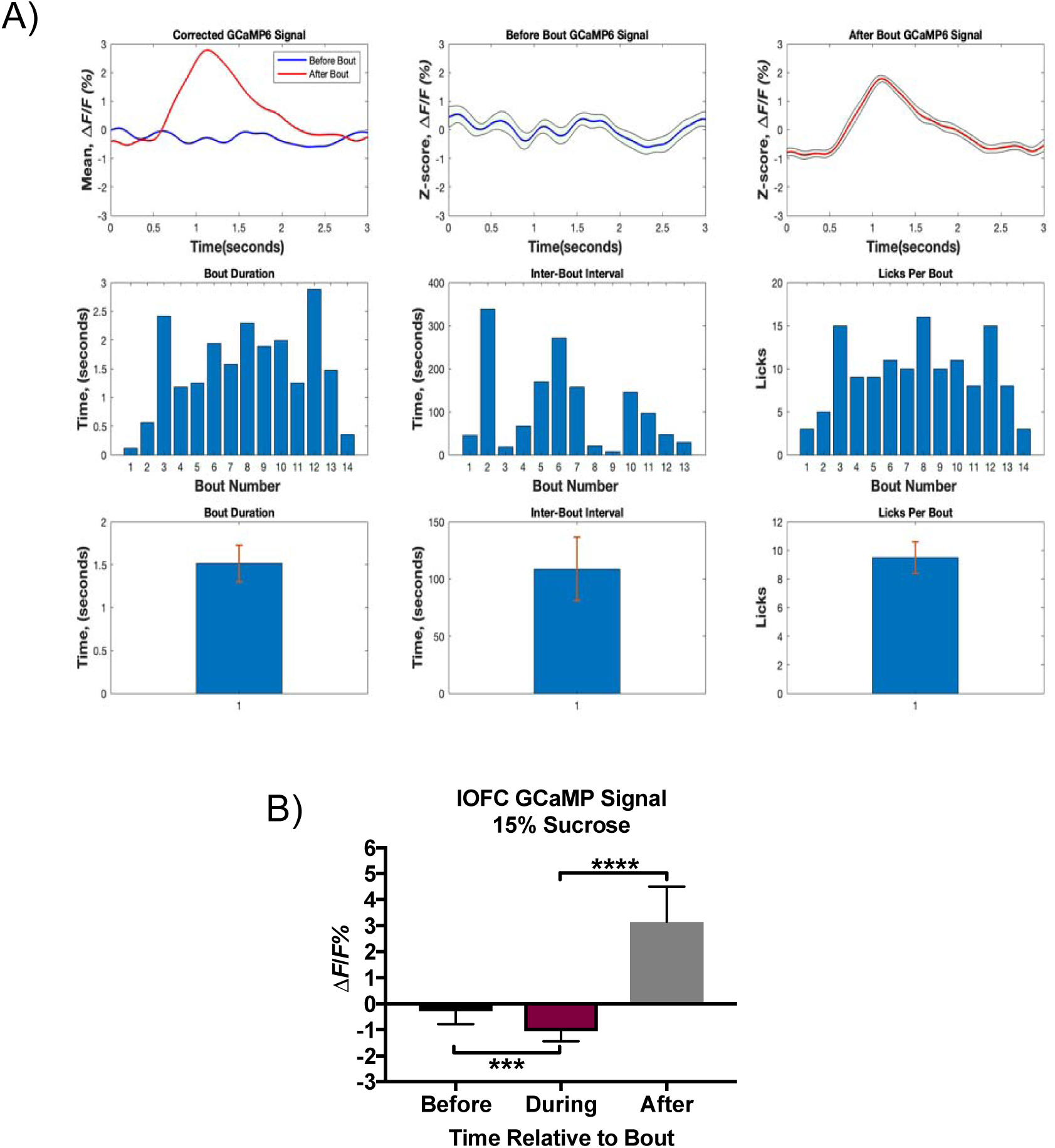
Example of activity of lateral OFC neurons in a C57BL/6J mouse during sucrose self-administration. (A) Top panels show mean and Z-scored ΔF/F values during bouts of drinking 15% sucrose. Middle panels show analysis of bout structure with bout duration (left), inter-bout interval (middle) and licks per bout (right) with summary plots (mean ± sem) below. (B) Mean (±sem) change in ΔF/F signal before, during and after bout. Symbol (***,****), p < 0.001, 0.0001, 1-way ANOVA.

Figure 4 summarizes data from studies of OFC activity using mice trained to self-administer 10% ethanol and then exposed to four weekly cycles of chronic intermittent ethanol (CIE) exposure that produces dependence and enhanced levels of homecage ethanol drinking and operant ethanol self-administration (den Hartog *et al*, 2016; Lopez and Becker, 2005). Prior to CIE, animals showed consistent lever pressing for ethanol (Fig. 4A) and like that seen for sucrose, OFC activity decreased during bouts of ethanol drinking followed by a rebound in signal when drinking stopped (Fig. 4B, C). After 4 weekly cycles of air or CIE exposure, animals underwent additional operant self-administration sessions for ethanol, ethanol plus quinine (60 µM) and ethanol plus sucrose (5%). Compared to initial levels of activity, both air and CIE treated mice showed similar decreases in signal during ethanol drinking and control-like increases in activity after the bout (Fig. 4E, F). When quinine was added to the ethanol solution, both air and CIE treated mice showed a greater decrease in ΔF/F during the bout as compared to ethanol only bouts while the post-bout spike in activity was similar to control values. During the ethanol + sucrose sessions, air-treated mice showed less reduction in signal during the bout and a smaller increase after the bout as compared to ethanol only drinking. In contrast, CIE treated mice had a greater decrease in ΔF/F during ethanol + sucrose drinking but no difference in the rebound of activity following the bout as compared to ethanol alone. The differential responses of Air and CIE treated mice to the ethanol + sucrose solution occurred despite similar amounts of responding under that condition (Fig. 4D). Although preliminary, these data suggest that CIE treatment may alter the OFC’s ability to discriminate between aversive and rewarding conditions thus contributing to heightened levels of ethanol intake.

**Figure 4.**
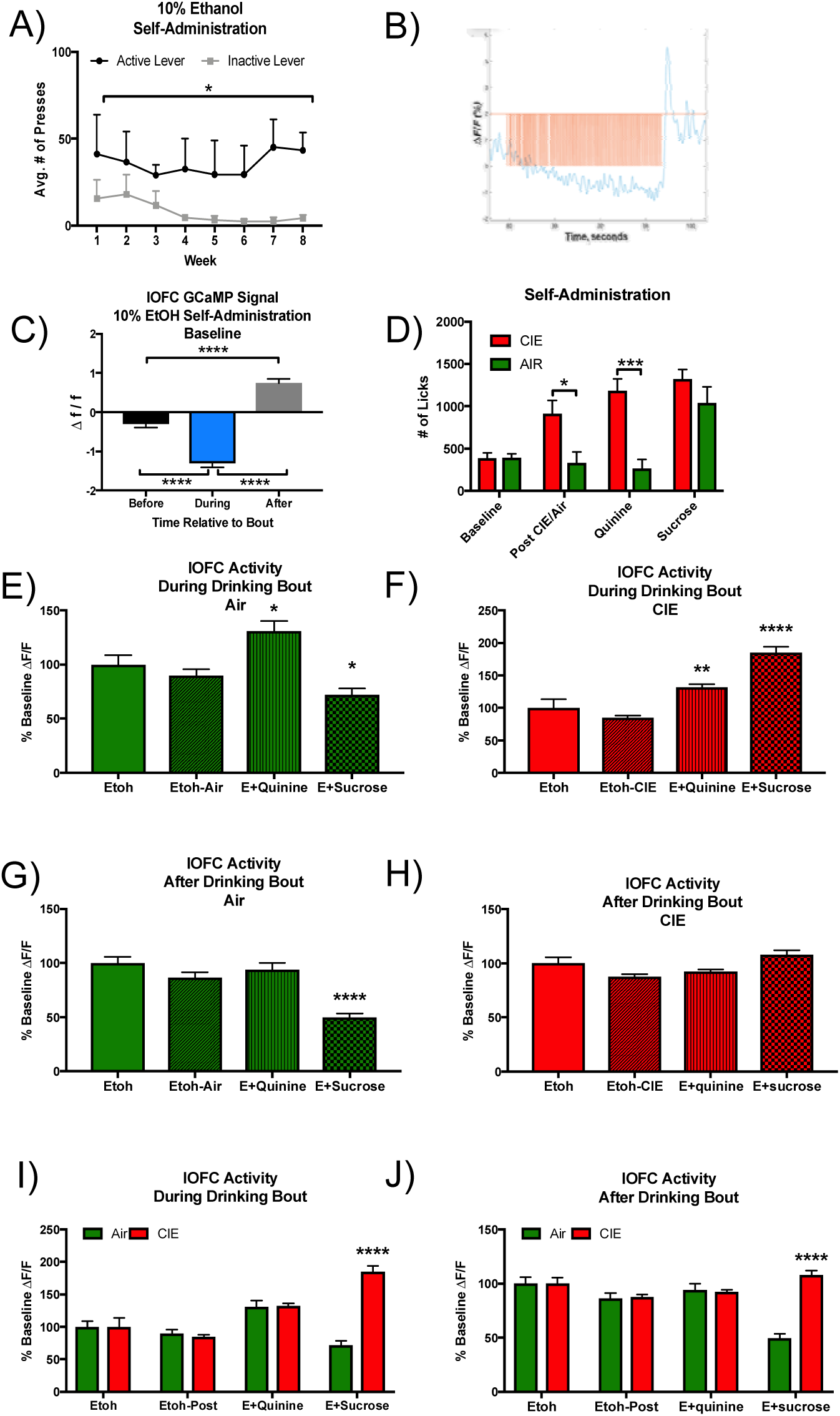
Activity of lateral OFC neurons in air and CIE-treated C57BL/6J mice during self-administration of ethanol, ethanol plus quinine or ethanol plus sucrose. (A) Mean ± sem presses on active and inactive levers prior to CIE exposure. (B) Example of GCaMP6 signal during ethanol drinking bout. Orange tick marks indicate licks. (C) Summary graph showing ΔF/F (mean ± sem) before, during and after ethanol drinking bouts in mice prior to CIE. (D) Lickometer data for the different drinking solutions in air and CIE exposed mice. All solutions contained 10% ethanol or ethanol plus quinine (60 µM) or sucrose (5%) where indicated. Graphs show ΔF/F (as % ethanol only baseline) in Air and CIE treated mice during (E, F) and after (G, H) bouts of ethanol, ethanol plus quinine or ethanol plus sucrose self-administration. (I, J) Summary of ΔF/F drinking data for Air and CIE mice showing pairwise comparisons. Symbols: (*,**,***,****), p < 0.05, 0.01, 0.001, 0.0001. 1 and 2-way Anova.

Overall, the results from these ongoing studies indicate the ability of in vivo fiber photometry to detect changes in neural activity in key nodes of the addiction neurocircuitry during exposures to different classes of abused substances. These changes occurred in response to behaviorally relevant concentrations of abused drugs and were time-locked to discrete epochs in the animal’s behavior. Importantly, neural activity in individual animals and the signature surrounding drinking behaviors was consistent over long periods of time allowing us to test a variety of conditions in the same animal. In addition, the photometry recordings identified ramping of activity in mPFC neurons consistent with results from in vivo electrophysiological studies (Horst *et al*, 2013) and changes in OFC signaling in ethanol-dependent mice. Together, these findings demonstrate the utility of this technology to capture physiologically-relevant neural events surrounding discrete behaviors that are important for the addiction process. Findings from this study add to a growing literature that fiber photometry is a powerful approach to record from principal output neurons in cortex and accumbens in rodent models of drug and alcohol addiction. Now that this approach is firmly established, future studies can apply advanced technologies to examine activity within specific circuits, presynaptic terminals, or subpopulations of cells within heterogeneous brain regions (e.g., prefrontal cortex, amygdala, nucleus accumbens) during drug-seeking behaviors.

## Acknowledgements

This work was supported by grants R37AA009986, R01DA013951 and P50AA01761 to JJW, T32AA007474, F32AA0226774 (DG), K01AA025110 (JAR), F31DA045485-01-A1 (KB), F32 DA042518-01-A1 (WW) and U01AA020930 and R01AA023288 to PJM.

